# 3D Centroidnet: Nuclei Centroid Detection with Vector Flow Voting

**DOI:** 10.1101/2022.07.21.500996

**Authors:** Liming Wu, Alain Chen, Paul Salama, Kenneth W. Dunn, Edward J. Delp

## Abstract

Automated microscope systems are increasingly used to collect large-scale 3D image volumes of biological tissues. Since cell boundaries are seldom delineated in these images, detection of nuclei is a critical step for identifying and analyzing individual cells. Due to the large intra-class variability in nuclei morphology and the difficulty of generating ground truth annotations, accurate nuclei detection remains a challenging task. We propose a 3D nuclei centroid detection method by estimating the “vector flow” volume where each voxel represents a 3D vector pointing to its nearest nuclei centroid in the corresponding microscopy volume. We then use a voting mechanism to estimate the 3D nuclei centroids from the “vector flow” volume. Our system is trained on synthetic microscopy volumes and tested on real microscopy volumes. The evaluation results indicate our method outperforms other methods both visually and quantitatively.

## 1. INTRODUCTION

Fluorescence microscopy is an important tool for imaging biological tissues in three dimensions [1]. nuclei detection is an essential step for identifying cells for quantitative analysis. For computer-aided diagnosis such as cell tracking and cell counting, it is frequently unnecessary to accurately delineate the boundaries of nuclei, only to accurately identify the centroid of each nucleus. However, due to the large volume size, variant nuclei morphology and intensity inhomogeneity, manually identifying the nuclei centroids in a 3D microscopy volume is so labor intensive that it is impractical. Thus, automated nuclei detection is necessary.

A segmentation-free approach described in [1] employs multiscale cube filtering to locally enhance pre-processed images and uses local maxima regions to determine candidate centroids without knowing the boundary of nuclei. In [2] a method that uses gradient flow tracking and local adaptive thresholding for dense nuclei segmentation is described. Similarly, a method known as active contours or “snakes” [3] segments the object by iteratively minimizing the energy function and actively adjusting the contour shape to fit the object. The image analysis tool CellProfiler [4] provides customized image segmentation modules for identifying and quantifying cell phenotypes. More recently, Volumetric Tissue Exploration and Analysis (VTEA) [5], a toolkit in ImageJ, has been developed to efficiently segment nuclei and quantitatively analyze large tissue volumes. These methods described above generally suffer from the effects of noise and intensity inhomogeneity of the microscopy imaging acquisition process. This has been addressed in [6], which uses 3D active contours with inhomogeneity correction for segmenting low contrast fluorescence microscopy images.

More recently, machine learning and in particular Convolutional Neural Networks (CNNs) have provided a different approach for nuclei detection. The U-Net [7], a popular network for semantic image segmentation, uses an encoder-decoder architecture with shortcut concatenation for segmenting biomedical images. A modified U-Net has been used in [8, 9, 10] for nuclei segmentation in 3D fluorescence microscopy images. For these semantic segmentation models, post-processing such as watershed is typically required for identifying different nuclei centroids [11]. This has been improved in [12, 13] by using a vector field energy gradient map for detecting nuclei centroids and boundaries. These two methods can only work on 2D images and use large amounts of hand annotated images for training. Alternatively, [14] works on 3D volumes by fusing and reconstructing 2D segmentation results of each slice in a volume from three directions into a 3D segmentation mask. However, using a 2D to 3D reconstruction may not fully capture the 3D information of the volume. Another deep learning approach for detection and segmentation is Regional Convolutional Neural Networks (R-CNN). The use of R-CNN for nuclei detection in microscopy images was described in [15]. However, these models are designed for object detection in 2D images. This was addressed in [16], where a slice-and-cluster strategy combined with Faster R-CNN and hierarchical clustering was proposed to estimate the true nuclei centroids in a 3D fluorescence microscopy volume.

The deep learning-based methods described above typically rely on large amounts of training samples to achieve good performance and avoid over-fitting [17]. Manually annotating large numbers of nuclei masks in 3D microscopy volumes is labor-intensive. Data augmentation techniques are necessary for generating more training data. Traditional data augmentation approaches use random transformation and deformation techniques to create images but cannot significantly improve the performance of the network and are vulnerable to adversarial attacks [18]. More recently, deep learning-based techniques for data augmentation use Generative Adversarial Networks (GANs) to generate high quality realistic images [19]. In [8], a Spatially Constrained Cycle-Consistent Adversarial Network (SpCycleGAN) was proposed to map synthetic binary ellipsoidal nuclei masks to synthetic microscopy images. Similarly, [20] uses Bézier curves and an SpCycleGAN to generate non-ellipsoidal shaped nuclei.

In this paper, we propose a 3D CentroidNet for 3D nuclei centroid detection, which is an extension of the 2D method described in [12]. Our network directly works on 3D volumes and automatically generates synthetic volumes for training. The nuclei centroids are estimated using a robust vector flow voting mechanism, which is achieved by collecting votes from each voxel and followed by non-maximum suppression. These voxels are constructed as 3D vectors that point to their nearest nuclei centroids in a 3D vector flow volume. To address the data limitation issue, we train our model on synthetic microscopy volumes generated from an SpCycleGAN and evaluate on three different types of microscopy volumes.

## 2. PROPOSED METHOD

Our proposed 3D nuclei centroid detection method is shown in Figure 1. The system consists of two components!: (1) 3D synthetic data generation, and (2) 3D CentroidNet training and inference.

**Fig. 1.**
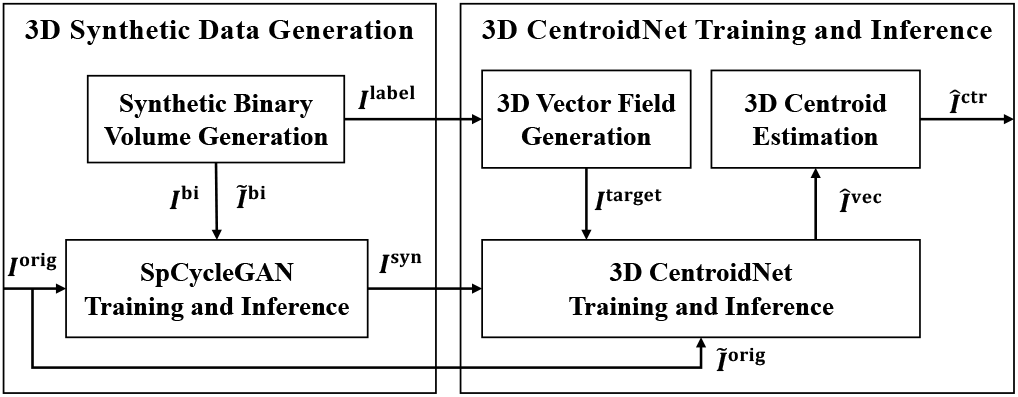
The block diagram of the proposed method

In this paper, we denote *I* as a 3D volume of size *X × Y × Z*. *I*(*χ,y,z*) denotes a voxel of I at location (*x,y,z*). Superscripts will be used to denote the type of volume. For example, *I*^orig^ and *I*^syn^ denote the original 3D microscopy volumes and the synthetic 3D microscopy volumes generated from SpCycleGAN. *I*^bi^ and *ℨ*^bi^ denote different synthetic 3D binary volumes for training and inference SpCycleGAN, and *I*^label^ is the corresponding label of the nuclei where each nucleus is marked with a unique voxel intensity. We define “vector flow” as a 3-channel volume where each voxel location is a 3D vector that points to its nearest nuclei centroid in the corresponding microscopy volume.

The SpCycleGAN was trained on *I*^bi^ and *I*^orig^ and inferenced on *ℨ*^bi^ to generate *I*^syn^ [8]. The pairs of *I*^syn^ and *ℨ*^bi^ will be further used for training a 3D CentroidNet. *I*^target^ is a four-channel 3D volume generated from *I*^label^. The first three channels correspond to the nuclei centroid location where each voxel represents a 3D vector 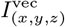 and points to its nearest nucleus centroid. The details of 3D vector flow generation and 3D centroid estimation will be discussed in Section 2.2. The last channel is the binary mask volume *ℨ*^bi^ which is used for segmentation. *I*^target^ serves as the ground truth of *I*^syn^ for training the 3D CentroidNet. The output of 3D CentroidNet consists of two parts: *Î*^vec^ and *Î*^mask^, where *Î*^vec^ is a three-channel volume which represents the estimated 3D vector flow, and *Î*^mask^ is the segmentation mask. *Î*^ctr^ is a 3D binary volume where the highlighted voxels represent the nuclei centroids.

### 2.1. 3D Synthetic Data Generation

We use synthetic microscopy volumes for training the proposed 3D CentroidNet. The synthetic data generation involves two steps: (1) Synthetic binary volume generation, and (2) SpCycleGAN training and inference.

#### Synthetic binary volume generation

We first generate synthetic 3D binary volumes *I*^bi^ and corresponding label volume *I*^label^ where each nucleus is corresponding to a unique gray-scale intensity. With the assumption that nuclei are ellipsoidal, we iteratively generate *N* ellipsoidal candidate nuclei having different sizes and orientations, where N is selected based on the nuclei density in *I*^orig^. The size is defined by the semi-axis length **a** = (*a_x_,a_y_,a_z_*) of an ellipsoid and the orientation is defined by a random rotation based on the translation matrix described in [21]. The k^th^ candidate nuclei *I*^can,k^ is generated with voxel intensity *k* ∈ {1,..., *N*} and added to *I*^label^ in random locations.

#### Synthetic microscopy volume generation

The synthetic microscopy volume *I*^syn^ is generated from the Spatially Constrained CycleGAN (SpCycleGAN) [8] trained with unpaired synthetic binary volumes *I*^bi^ and the original microscopy volumes *I*^orig^. As shown in Figure 2, SpCycleGAN consists of 5 networks *G, F, H, D*_1_, and *D*_2_. *G* and *F* are two generators that map an image from one domain to another. *D*_1_ and *D*_2_ are discriminators for two domains to distinguish if the given images are real or synthetic. *H* is the extra generator identical to *F* for maintaining the spatial alignment between a microscopy volume and its corresponding mask. The entire loss function of SpCycleGAN *L*_1_ is shown in Equation 1,

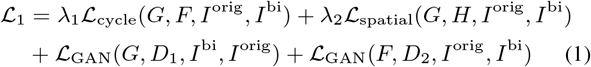

where 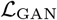 is the discriminator loss, and λ_1_, λ_2_ are weight coefficients controlling the loss balance between the cycle consistency loss 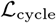 and the spatial constraint loss 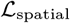 given by a *L*_2_ norm,

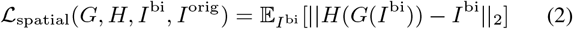

**Fig. 2.**
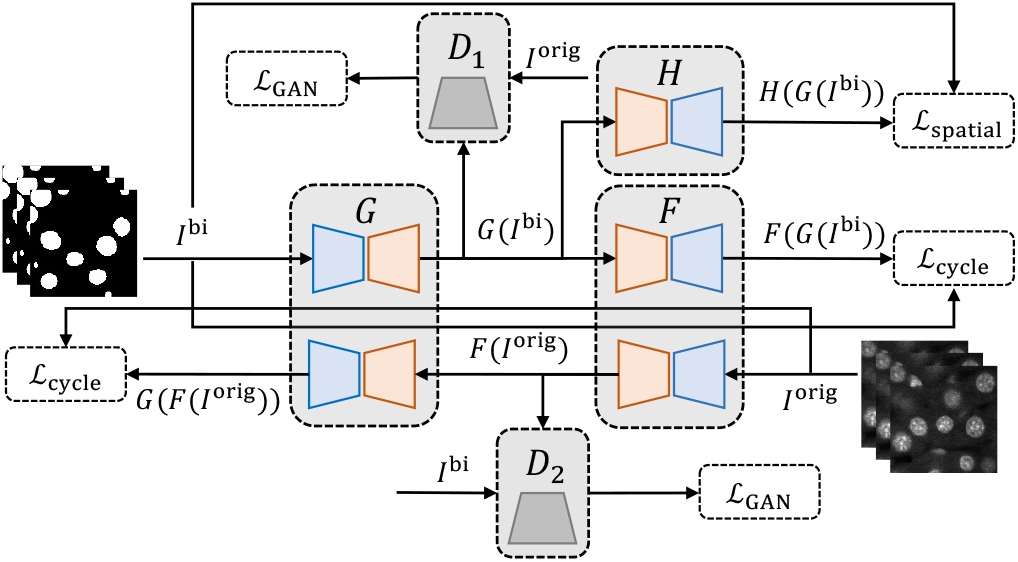
Architecture of SpCycleGAN.

### 2.2. 3D CentroidNet

#### Vector flow generation

The 3D vector flow generation prepares the target volumes *I*^target^ for training the 3D CentroidNet. It takes *I*^label^ as the input, where each nucleus is marked by a unique voxel intensity, and generates the centroid of each nucleus. Then the volume *I*^vec^ is generated, where each voxel (*x,y,z*) has 3 channels that represent a 3D vector 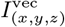 that points to its nearest nucleus centroid. The voting region of a nucleus centroid (shown within the dashed circles in Figure 3) is determined by a radius threshold *T*_vec_. The voxels outside the voting region are set to zero and the voxels inside the voting region are encoded with a 3D vector pointing to the nearest nucleus centroid (the center of the circle). The area of the voting region is set to the same size for each nucleus so each voxel can maximally collect an equal number of votes. *I*^target^ is the concatenation of *I*^vec^ and *I*^bi^ used during training 3D CentroidNet.

**Fig. 3.**
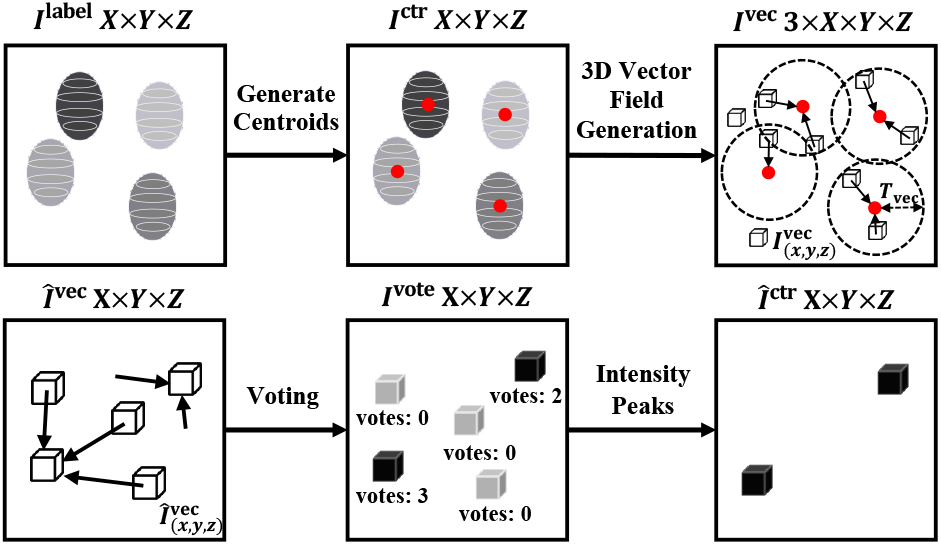
Overview of 3D vector flow generation (first row) and 3D centroid estimation (second row)

#### 3D CentroidNet

To learn the nuclei centroid location using our proposed vector flow voting mechanism, we propose a network known as 3D CentroidNet shown in Figure 4 that consists of a head module and a backbone network. The “head” is a multi-task learning module for learning the vector flow volume and performs vector flow voting to estimate nuclei centroids. The “backbone” is simply a modified 3D UNet [7]. We use an encoder-decoder network with shortcut concatenation as the backbone so it can take a volume in any dimension and output a volume with identical size. The encoder consists of multiple convolution blocks and 3D max pooling layers. Each convolution block consists of a 3D convolution layer with filter size 3 × 3 × 3, a 3D batch normalization layer, and a leaky ReLU layer. For the decoder, conversely, each 3D transpose convolution block consists of a 3D transpose convolution layer followed by a 3D batch normalization and a leaky ReLU layer.

**Fig. 4.**
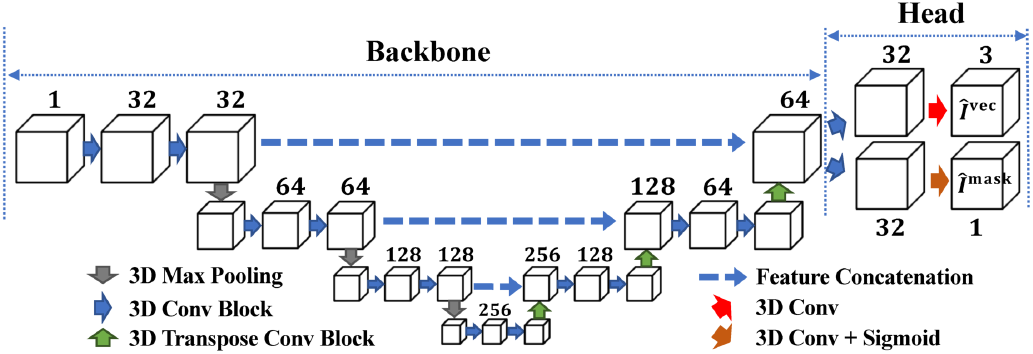
The architecture of our proposed 3D CentroidNet.

#### Multitask learning

The head of the network consists of two branches. One branch outputs the estimated vector flow volume *Î*^vec^, and another branch outputs the binary segmentation masks *Î*^mask^. The segmentation branch can help the network identify the boundary of nuclei and converge faster due to the extra loss constraint. Note that there is no sigmoid function used to obtain *Î*^vec^ because the voxel values can be a negative number or a large number. In other words, the vector encoded in a voxel can point to anywhere in a volume. During training, the synthetic volumes *I*^syn^ generated from SpCycleGAN are used as the input, and the output volumes from two branches *Î*^mask^ and *Î*^vec^ are compared with *I*^target^ for optimization. Specifically, the output vector flow volumes *Î^vec^* are compared with the ground truth vector flow volumes *I*^vec^ and optimized using the Mean Square Error (MSE) loss function, whereas the segmentation results *Î*^mask^ are compared with the ground truth binary volumes *Ĩ*^bi^ and optimized using the combination of Focal Loss [22] 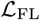 and Tversky Loss [23] 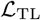. The total loss of our network 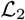 is shown in Equation 3.

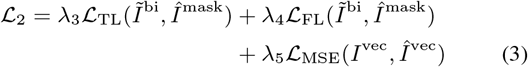

#### 3D centroid estimation

The steps for 3D centroid estimation are shown in Figure 3. *Î*^vec^ is a 3-channel volume where each voxel is a 3D vector that points to somewhere in the volume. We denote the “votes” of a voxel 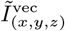 as the number of other voxels that point to it. For example, if there are 3 different vectors that point to 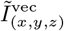, then the voxel 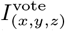 will have 3 votes. A voting map volume *I*^vote^ is generated where the intensity of each voxel is the number of votes for that voxel to be a centroid. Thus, the higher voxel intensity in a voting map indicates higher probability of being a centroid. Nonmaximum suppression with window threshold size *T*_ctr_ is used to remove duplicate centroids. Finally, we use intensity thresholding to remove the voxels with votes less than *T*_vote_. The details of the parameter values are shown in Table 1. The visualization of the 3D vector flow is shown in Figure 5.

**Table 1.** Parameters for synthetic binary volume generation, 3D centroid estimation, and object centroid-based evaluation

| | $r_{min}$ | $r_{max}$ | $T_{ov}$ | $N$ | $T_{vec}$ | $T_{ctr}$ | $T_{vote}$ | $T_{match}$ |
| --- | --- | --- | --- | --- | --- | --- | --- | --- |
| Data-I | 4 | 8 | 5 | 400 | 10 | 7 | 30 | 4 – 8 |
| Data-II | 10 | 14 | 10 | 40 | 24 | 7 | 200 | 6 – 12 |
| Data-III | 6 | 12 | 20 | 200 | 16 | 7 | 70 | 6 – 12 |

**Fig. 5.**
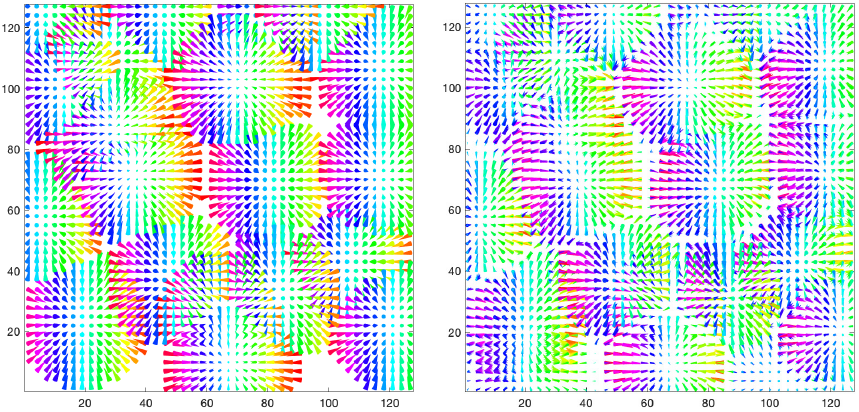
Visualization of the ground truth vector flow volume (left) and the estimated 3D vector flow volume (right) for a volume in real microscopy Data-II. Each cone represents a 3D vector pointing to its nearest nuclei centroid

## 3. EXPERIMENTAL RESULTS

### 3.1. Experimental Setup

Our proposed method is trained on synthetic microscopy volumes and tested on three different real microscopy data denoted as Data-I, Data-II and Data-III. Data-I is a gray-scale volume of size *X × Y × Z =* 128 × 128 × 64 voxels, whereas Data-II includes 16 volumes of size 128 × 128 × 16 voxels and Data-III consists of 4 volumes of size 128 × 128 × 64 voxels. These data are collected from rat kidney using two-photon microscopy and the ground truth volumes are manually annotated using ITK-SNAP [25]. The original Data-I, Data-II, and Data-III can be obtained from [9]. For corresponding synthetic volumes, the parameters for generating *I*^bi^ are appropriately chosen based on the nuclei size in real microscopy volumes. As shown in Table 1, rmin and rmax are the minimum and maximum semi-axis length of the nuclei size. Tov is the maximum allowed overlapping voxels between two nuclei. *N* is the total number of nuclei in a synthetic volume. The SpCycleGAN is trained on 4 pairs of *I*^bi^ and *I*^orig^ individually for each data with λ_1_ = λ_2_ = 10, and inferenced on 50 other synthetic binary volumes *Ĩ*^bi^ to generate synthetic microscopy volumes *I*^syn^ for training 3D CentroidNet.

For 3D CentroidNet, 3 different models denoted as M-I, M-II, and M-III are trained on synthetic Data-I, Data-II, and Data-III, respectively. Since Data-III consists of both ellipsoidal and non-ellipsoidal nuclei whereas our synthetic data generated from SpCycleGAN contains only ellipsoidal nuclei, we observed that M-III does not generalize well on real microscopy Data-III. Thus, we updated M-III on 3 volumes of real Data-III with the same parameter for training synthetic Data-III. The 3D CentroidNet was implemented in PyTorch and trained for 100 epochs with the Adam optimizer and initial learning rate 0.001. The weight coefficients of the loss function is set to λ3 = 1, λ_4_ = 10, λ_5_ = 10. As shown in Table 1, the parameters for 3D centroid estimation are chosen based on the average nuclei size in the volume.

### 3.2. Quantitative Evaluation

To quantitatively evaluate the centroid detection accuracy of our model, we use object centroid-based precision 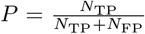 recall 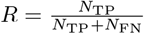, and 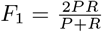 scores to evaluate the centroid detection accuracy. For each detected centroid, we define that it is “matched” with a ground truth centroid if their Euclidean distance is less than a centroid distance threshold *T*_match_. Note that each ground truth centroid always matches with its nearest detected centroid if there are multiple detections. *N*_TP_ is the number of True Positives (“matched” pairs). Similarly, *N*_FP_ is the number of False Positives (number of detected centroids with no associated ground truth centroid matched), and *N*_FN_ is the number of False Negatives (number of remaining no matched ground truth centroids). *T*_match_ is chosen based on the inspection of real microscopy data and verified by a biologist. The details of *T*_match_ is shown in Table 1. In addition, we use the Average Precision (AP) and mean Average Precision (mAP) that was presented in [16] for object centroid-based evaluation. AP_*t*_ is the AP score with the distance threshold *T*_match_ = *t*. The performance of our method was compared with VTEA [5], CellProfiler [4], Squassh [26], and three deep learning methods DeepSynth [9], Cellpose [14], and RCNN-SliceNet [16]. We use Cellpose to first obtain the 3D instance segmentation masks. We then extract the centroids for each nucleus obtained from Cellpose. The RCNN-SliceNet is capable of capturing almost all nuclei but suffers from over-detection, whereas DeepSynth missed some non-ellipsoidal nuclei. Our approach uses a one stage method and addresses the limitations of RCNN-SliceNet which requires a rough estimation of the nuclei number. Also, since our method works directly on 3D volumes using a 3D CNN, it can make better use of the 3D information of a volume. The object centroid-based evaluation results shown in Figure 6, Table 2 and Figure 7 indicate that our method achieved best visual performance as well as quantitative accuracy towards mAP and *F*_1_ scores on three different microscopy data.

**Fig. 6.**
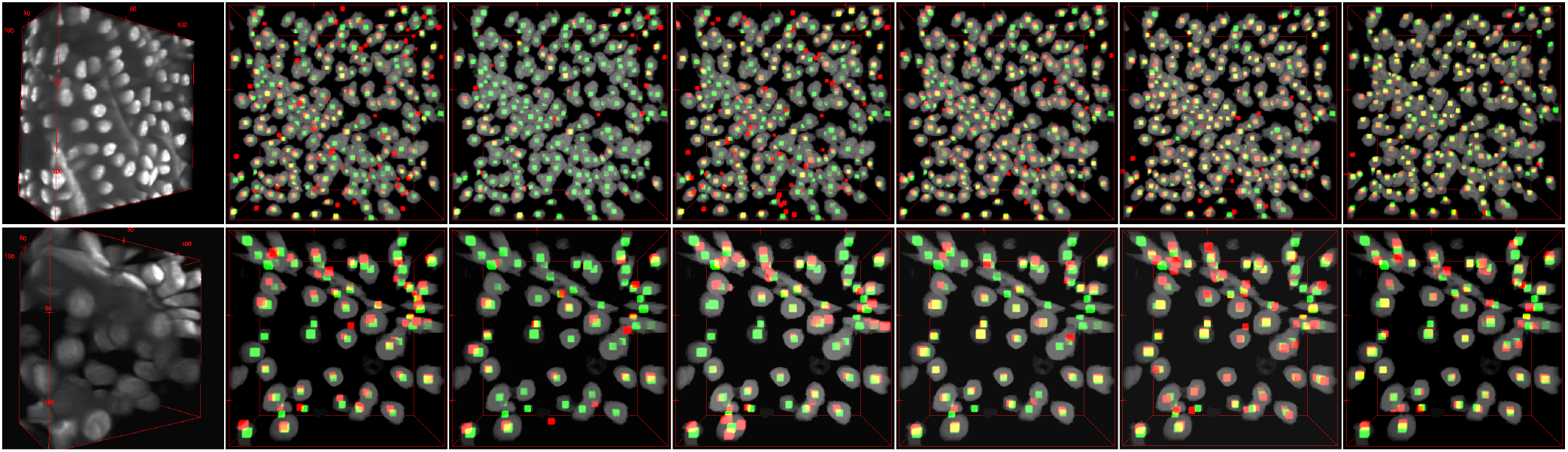
Visual evaluation using ImageJ’s 3D visualization tool [24]. The first column are example testing volumes from original microscopy Data-I and Data-III. The nuclei centroid estimation results for CellProfiler (second column), Squassh (third column), VTEA (fourth column), DeepSynth (fifth column), RCNN-SliceNet (sixth column), and our 3D CentroidNet (last column). The estimated centroids are red and the ground truth centroids are green

**Fig. 7.**
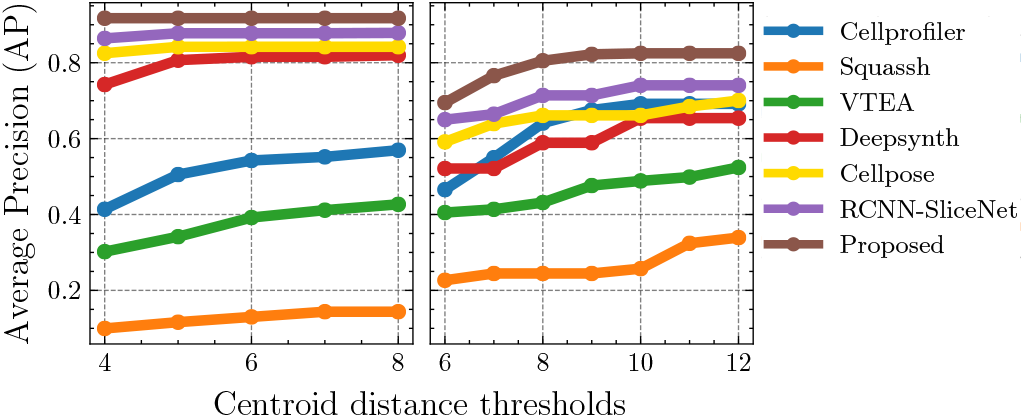
Comparison of the object centroid-based evaluation results using Average Precision (AP) for Data-I (left) and Data-III (right)

**Table 2.**
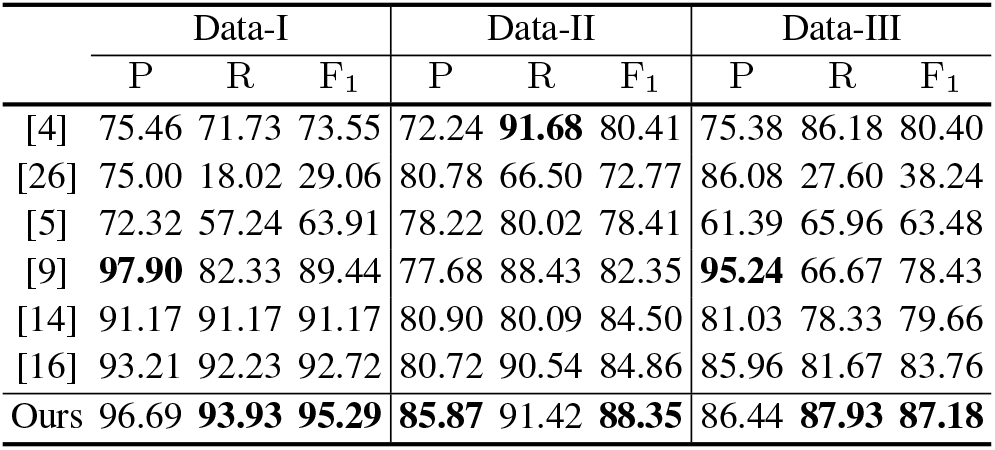
Comparison of the object centroid-based evaluation results using Precision (P), Recall (R), and F_1_ scores.

|  | Data-I |  |  | Data-II |  |  | Data-III |  |  |
| --- | --- | --- | --- | --- | --- | --- | --- | --- | --- |
|  | P | R | F<sub>1</sub> | P | R | F<sub>1</sub> | P | R | F<sub>1</sub> |
| [4] | 75.46 | 71.73 | 73.55 | 72.24 | **91.68** | 80.41 | 75.38 | 86.18 | 80.40 |
| [26] | 75.00 | 18.02 | 29.06 | 80.78 | 66.50 | 72.77 | 86.08 | 27.60 | 38.24 |
| [5] | 72.32 | 57.24 | 63.91 | 78.22 | 80.02 | 78.41 | 61.39 | 65.96 | 63.48 |
| [9] | **97.90** | 82.33 | 89.44 | 77.68 | 88.43 | 82.35 | **95.24** | 66.67 | 78.43 |
| [14] | 91.17 | 91.17 | 91.17 | 80.90 | 80.09 | 84.50 | 81.03 | 78.33 | 79.66 |
| [16] | 93.21 | 92.23 | 92.72 | 80.72 | 90.54 | 84.86 | 85.96 | 81.67 | 83.76 |
| Ours | 96.69 | **93.93** | **95.29** | **85.87** | 91.42 | **88.35** | 86.44 | **87.93** | **87.18** |

## 4. CONCLUSION

In this paper, we proposed an approach, 3D CentroidNet, to detect the centroid of nuclei in 3D microscopy volumes. The nuclei centroid detection is achieved by a robust vector flow voting mechanism. We demonstrate that 3D CentroidNet outperforms the compared methods in nuclei centroid detection on three microscopy datasets. 3D CentroidNet is able to work on any size of input volumes. Our method is practical to use since the user only needs to provide original microscopy volumes and our network will automatically generate synthetic microscopy volumes for training 3D CentroidNet and provide 3D detections for these original microscopy volumes. The code and dataset are available upon request to

